# The effects of YKL-40 on angiogenic potential of HUVECs are partly mediated by syndecan-4

**DOI:** 10.1101/2020.04.19.049064

**Authors:** Jianlei Zheng, Qi Xue, Yan Zhao, WeiJun Sun

## Abstract

**Objective:** YKL-40, a secreted glycoprotein, has a role in promoting tumor angiogenesis through syndecan-1 receptor. As one of the members of syndecans family, syndecan-4 is also an important mediator for tube formation. However, the effects of YKL-40 on migration and tube formation of human umbilical vein cells (HUVECs) mediated by syndecan-4 receptor are unknown.

**Methods:** HUVECs were transfected with lentivirus encoding syndecan-4 short hairpin (sh) RNA (lenti-synd4 shRNA) and the efficiency of transfection was measured using reverse transcription quantitative real time polymerase chain reaction (qRT-PCR) and flow cytometry. The effects of recombinant protein of YKL-40 on migration and angiogenesis of HUVECs adjusted by syndecan-4 were determined by wound healing and tube formation assay. The expression of protein kinase Cα (PKCα) and extracellular signal regulated kinases (ERKs) 1 and 2 (ERK1/2) in HUVECs was measured using western blotting.

**Results:** Transfection of lenti-synd4 shRNA significantly decreased the membrane protein expression of syndecan-4 in HUVECs. HUVECs transfected with lenti-synd4 shRNA remarkably inhibited the migration and tube formation of endothelial cells stimulated by recombinant protein of YKL-40. The levels of PKCα and ratio of p-ERK1/2 to ERK1/2 in HUVECs were also down-regulated by silencing syndecan-4.

**Conclusion:** The effects of YKL-40-induced on migration and tube formation of HUVECs may partly be inhibited by knock-downing syndecan-4 through suppressing PKCα and ERK1/2 signaling pathways.

## Introduction

YKL-40 (also known as human cartilage glycoprotein-39) is a 40 kDa heparin- and chitin-binding glycoprotein, which has been known to play an important role in cell migration, proliferation and angiogenesis.^1,2^ Recently, angiogenesis is regarded as a main characteristic for vulnerable plaques.^3^ Immature neovessels may contribute to intraplaque hemorrhage and inflammatory cells infiltration, which are critical factors in atherosclerotic plaque progression and destabilization.^4^ In clinical practices, it has been reported that elevated serum YKL-40 was associated with acute coronary syndrome and cardiovascular morbidity and mortality. ^5-7^ Hence, It is supposed that increased serum YKL-40 as a biomarker for adverse prognosis in cardiovascular diseases may be related to its role in atherosclerotic angiogenesis.

Syndecans (SDCs) are a family of heparan sulfate proteoglycans, including four mammalian syndecans, from syndecan-1 to syndecan-4, which play a critical role in cell adhesion, migration, proliferation and angiogenesis.^8,9^ The physiological and pathological function of SDCs is fulfilled by interacting with a great number of ligands including insoluble matrix collagens, associated glycoproteins and interleukins. Moreover, syndecans can also affect cell function via interacting with its adjacent membrane receptors such as integrin and fibroblast growth factor receptor (FGFR).^10,11^ All syndecans are shown to carry similar molecular structure of heparan sulfate chains, and heparan sulfate is regarded as a physiological ligand for YKL-40.^9,12^ It was reported that YKL-40-induced tumor angiogenesis was dependent on the interaction via YKL-40 and membrane receptors syndecan-1 as well as integrin αvβ3.^13^ Different from the other three syndecans, syndecan-4 has a more widespread tissue distribution. However, whether the angiogenic potential of YKL-40 in HUVECs is regulated by syndecan-4 is unknown, so the present study is to explore the effects of YKL-40 on migration and tube formation of HUVECs and potential signal pathways mediated by syndecan-4.

## Methods

### Cell culture

HUVECs were purchased from Shanghai Institutes for Biological Sciences, Chinese Academy of Sciences (Shanghai, China). HUVECs were grown in Roswell Park Memorial Institute (RPMI) 1640 medium from Gibco (Carlsbad, CA, USA) supplemented with 10% fetal calf serum (FCS) and 1% penicillin-streptomycin solution (Invitrogen, Carlsbad, CA, USA) and were incubated at 37□ in a humidified atmosphere containing 5% CO2. HUVECs in the present study were divided into control (HUVECs without additional treatment), YKL-40 group (HUVECs stimulated with 100ng/ml YKL-40, Cat No: 11227-H08H, Sino Biological Inc. China), YKL-40+lenti-synd4 shRNA group (HUVECs transfected with lentivirus encoding syndecan-4 shRNA plus 100ng/ml YKL-40) and YKL-40+lenti-null group (HUVECs transfected with lentivirus carrying scramble shRNA plus 100ng/ml YKL-40).

### Silencing syndecan-4 gene in HUVECs

At first, we designed three couples of short hairpin RNAs (shRNAs) and then they were transfected into human embryonic kidney (HEK)293 cells. The shRNAs with most powerful inhibition to syndecan-4 expression was selected for subsequent experiments (decreasing about 60 percent expression of syndecan-4 relative to control at mRNA level). A lentiviral vector silencing syndecan-4 expression was constructed as below. Briefly, The sequences used for short hairpin RNAs (shRNAs) in this study are as follows: shRNA (targeting syndecan-4): 5’-GCAGGAATCTGATGACT TTGATTCAAGAGATCAAAGT CATCAGATTCCTGCTTTTTT-3’; shRNA (scramble): 5’-GCACCCAGTCCG CCCTGAGCAAATTCAAGAGATTTGCTCAGGGCGGACTGGGTGCTTTT T-3’, which were subsequently cloned into pLent-U6-GFP-Puro. Infectious viruses were produced by cotransfecting the lentiviral vector and packaging constructs into 293FT cells (Invitrogen, Carlsbad, CA, USA) using transfection reagent (BioRad, Hercules, CA, USA). Infectious lentivirus particles were harvested at 48 hours after transfection. The final virus titres were approximately 5.0×10^8^ TU/mL, and the whole process of lentivirus construction was supplied by Shandong ViGene Biosciences (Jinan, Shandong, China). HUVECs at passage 2 were successfully infected with the corresponding virus at a multiplicity of infection (MOI) of 50 confirmed by fluorescence microscope. At last, cells transfected with the lentiviruses were further selected for 48 hours in medium containing 2ug/mL puromycin (Sigma-Aldrich, St. Louis, MO, USA) until the final stable cell clones were harvested and verified for following studies.

### The effect of lenti-synd4 shRNA interference was verified by qRT-PCR and flow cytometry

The total RNA was extracted from stable transfectants of HUVECs with TRIzol Reagent from Thermo Fisher Scientific (Waltham, MA, USA), and the RNA concentration was measured at 260nm/280nm absorbance ratio. The total RNA was reverse-transcribed to cDNA using a high capacity cDNA Reverse Transcription Kit from GeneCopoeia (Maryland, USA). qRT-PCR was carried out to evaluate the syndecan-4 expression between lenti-synd4 shRNA and lenti-null group with SYBP Premix Ex Taq TM from Takara (Shiga, Japan) following the manufacturer’s instructions. The primers for syndecan-4 and glyceraldehyde-3-phosphate dehydrogenase (GAPDH) as below: Homo syndecan-4 Forward: 5’-GGACCTCCTAGAAGGCCGATA-3’, Homo syndecan-4 Reverse: 5’-AGGGCCGATCATGGAGTCTT-3’. Homo GAPDH Forward: 5’-CTCGCTTCGGCAGCACA-3’; Homo GAPDH Reverse : 5’-AACGCTTCACGAATTTGCGT-3’. In addition, HUVECs were successively stained with goat anti syndecan-4 (Cat No: AF2918-SP, R&D Company, USA) and rabbit anti-goat IgG secondary antibody (Cat No: A27018, Thermo Fisher Scientific, USA). Then the syndecan-4 expression on HUVECs was measured by flow cytometry with a BD Biosciences LSR II (BD Biosciences, CA, USA).

### Wound healing assay

HUVECs at 95% confluence were plated at a density of 3×10^5^ cells/well in 6-well plates and cultivated in RPMI 1640 medium without FCS at 37□ overnight. The monolayer of HUVECs was scratched with a sterile pipette tip to form wound gaps. Cells were then washed to remove debris by PBS and incubated with or without 100ng/ml YKL-40 at 37□ to induce cell migration for 24 hours and 32 hours. Images from two different scratch areas in each culture well were systematically obtained using a light microscope (magnification, ×100) equipped with Leica Application Suite program (Leica Microsystems, Switzerland). The distance between wound edges was measured at least 3 points on each image, and then the percent gap closure were calculated at 24 hours and 32 hours relative to that at 0 hour.

### Tube formation assay

Tube formation assay was performed to evaluate angiogenic ability of HUVECs. Briefly, the cells were cultured in medium containing 10% FCS with or without 100ng/ml YKL-40 for 24 hours and 48 hours, then were seeded into a 96-well plate at a density of 5×10^3^cells/well coated with 10mg/ml Matrigel matrix without growth factor (Cat No: 3432-010-01, BD Biosciences, CA, USA) for 6 hour incubation at 37□. Capillary tube structures were observed and representative images were captured using Leica Application Suite program (magnification, x100). The value of branch points × number of branches representing the capacity of angiogenesis was determined using Image J software.^14^

### Western blotting analysis

HUVECs transfected with lenti-synd4 shRNA or lenti-null grown in 60 mm dishes were treated with 100ng/ml YKL-40 for 24 hours and 48 hours. Cells were lysed using Radio-Immunoprecipitation assay (RIPA) lysis buffer from Thermo Fisher Scientific (Waltham, MA, USA) at 4°C for 30 minutes, and subjected to 12000 rpm centrifugation at 4°C for 5 minutes. The protein concentration was measured by BCA protein assay Kit from Thermo Fisher Scientific (Waltham, MA, USA). The lysates were separated by 12% sodium dodecyl sulfate-polyacrylamide gel electrophoresis (SDS-PAGE), then transferred to polyvinylidene fluoride (PVDF) membranes and blocked with 5% skim milk. The blots were analyzed with antibodies according to the manufacturer’s instructions and visualized by peroxidase and an enhanced chemiluminescence system from Pierce Biotechnology (Waltham, MA, USA). The corresponding antibodies were used in the present study including anti-p-ERK1/2 and anti-ERK1/2 (Cat No: YP1197, Immuno Way, USA; Cat No: ab65142, Abcam, Cambridge, UK, respectively); anti-PKCα and anti-GAPDH (Cat No: ab32376 and ab181602, Abcam, Cambridge, UK, respectively). Goat anti-rabbit IgG secondary antibody (Cat No: ab205718, Abcam, Cambridge, UK).

### Statistical analysis

Data analysis was performed with SPSS13.0 (SPSS, Inc., Chicago, IL, USA) software. Data were expressed as mean ± standard deviation. Comparisons between two groups were evaluated by t-test. A p-value less than 0.05 was considered statistically significant.

## Results

### Lenti-synd4 shRNA inhibited syndecan-4 expression in HUVECs

Fig.1A indicated that the HUVECs were successfully transfected with lentivirus confirmed by fluorescence microscope. To evaluate the inhibition effects of lenti-synd4 shRNA on syndecan-4 expression, we examined the gene and protein levels of syndecan-4 in HUVECs after successful viral transfection. The qRT-PCR test demonstrated that lenti-synd4 shRNA significantly inhibited the mRNA expression of syndecan-4 in lenti-synd4 shRNA group than that in lenti-null group (p<0.001), which was shown in Fig.1B. Flow cytometry was further performed to analyze the membrane receptor expression of syndecan-4 on HUVECs. The data from the Fig.1C-D showed the syndecan-4 expression on the cell membrane of HUVECs was also remarkably decreased in lenti-synd4 shRNA group, compared with the lenti-null group (p<0.001).

**Fig. 1.**
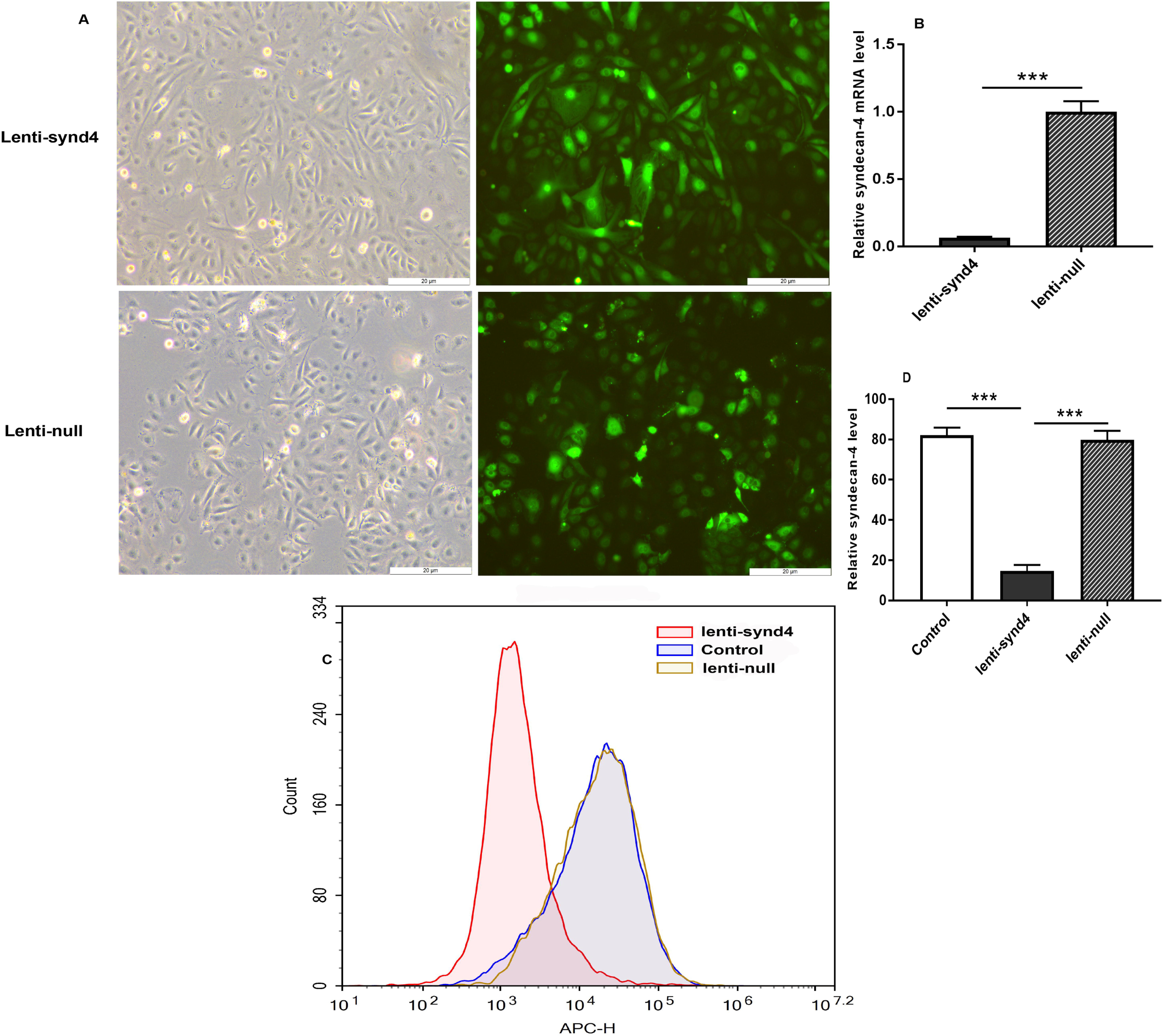
The mRNA and membrane protein expression of syndecan-4 from HUVECs after being transfected with lenti-synd4 shRNA. The lenti-synd4 shRNA and lenti-null shRNA were successfully transfected into HUVECs confirmed by fluorescence microscope (Fig.1A). The difference of mRNA production of syndecan-4 between HUVECs transfected with lenti-synd4 shRNA and lenti-null shRNA (Fig.1B). Flow cytometry analysis of membrane receptor of syndecan-4 on HUVECs was performed among control, lenti-synd4 shRNA and lenti-null group (Fig.1C and 1D). Data were presented as mean ± S.E. (n=3 per group).*** *p*<0.001.

### The effect of lenti-synd4 shRNA on the migration of HUVECs treated with YKL-40

Wound healing assays for evaluating migration of HUVECs were displayed in Fig.2A-C. We found that YKL-40 at concentration of 100ng/ml mildly prompted the migration of HUVECs compared with control at 24 hours (p<0.05), and the difference between 100ng/ml YKL-40 group and control were further extended at 32 hours (p<0.01). Meanwhile, lenti-synd4 shRNA group significantly decreased the migration rate of HUVECs stimulated with 100ng/ml YKL-40 after 24 hours and 32 hours (p<0.05 and p<0.01, respectively).

**Fig. 2.**
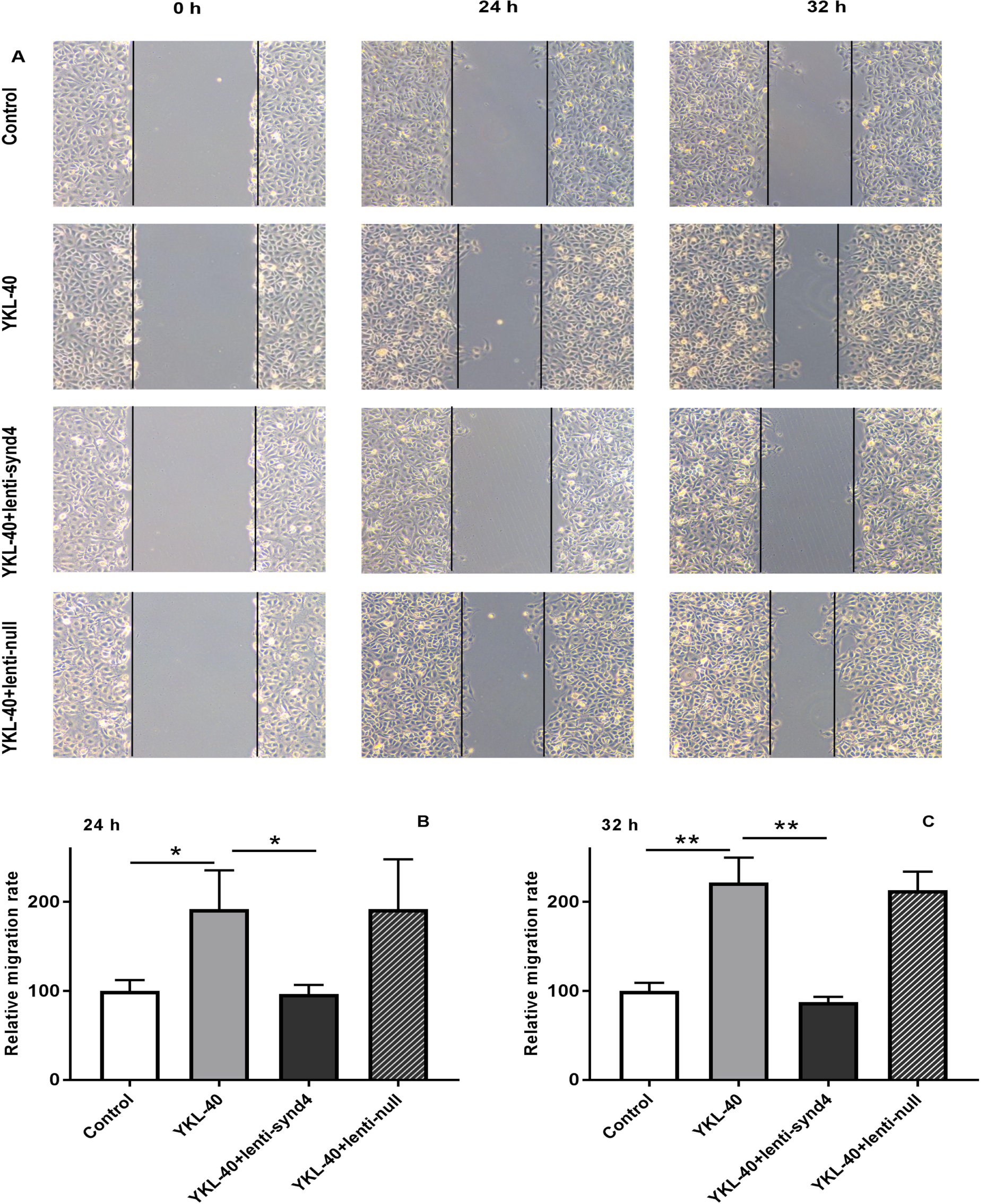
The effect of lenti-synd4 shRNA on the migration of HUVECs treated with YKL-40 at 24 hours and 32 hours. The representative photomicrographs of wound healing assay for control, YKL-40 group, YKL-40+lenti-synd4 shRNA group and YKL-40+ lenti-null group (Fig.2A). Photos were taken at 24 hours and 32 hours. The relative migration rate was calculated according to the ratio of initial width to terminal width and rectified by control (Fig.2B-C). Data were presented as mean ± S.E. (n=3 per group).** *p*<0.01 and \**p*<0.05.

### The effect of lenti-synd4 shRNA on the tube formation of HUVECs treated with YKL-40

YKL-40 was used to verify its effect on the tube formation in HUVECs. As shown in Fig. 3A-C, compared with the control, tube formation ability of HUVECs stimulated with 100ng/mL YKL-40 was remarkably increased at 24 hours and 48 hours (both p<0.01). We also found that HUVECs transfected with lenti-synd4 shRNA decreased the ability of tube formation mediated by YKL-40 at 24 hours (p<0.05). With the extension of culture time, compared with YKL-40 group, capacity of angiogenesis of HUVECs transfected with lenti-synd4 shRNA was significantly inhibited at 48 hours (p<0.01), which showed that the effects of YKL-40 on angiogenesis of HUVECs probably inhibited by lenti-synd4 shRNA were time-dependent manner.

**Fig. 3.**
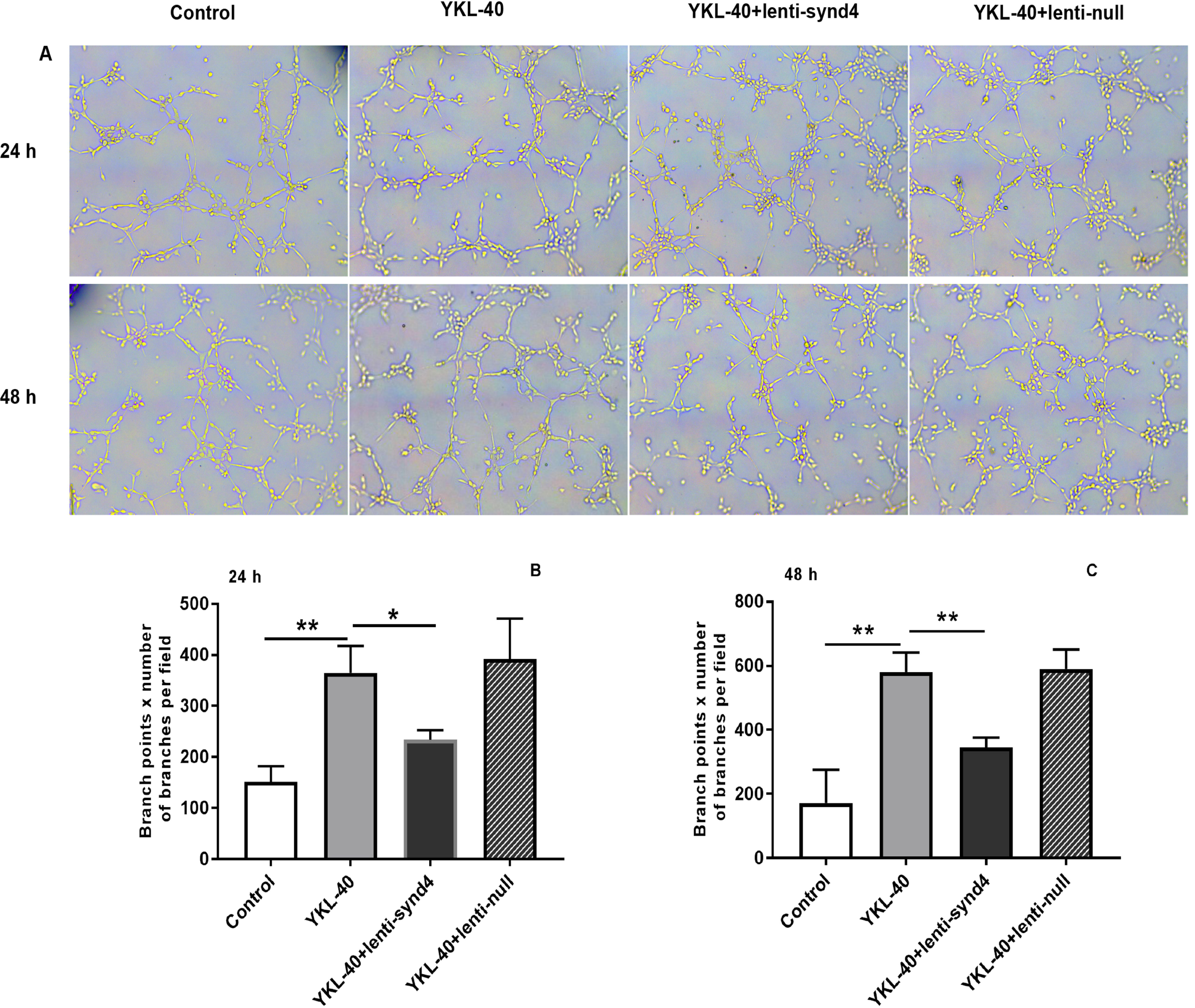
The effect of lenti-synd4 shRNA on the tube formation of HUVECs treated with YKL-40 at 24 hours and 48 hours. Representative microscopic images of the tube formation in different groups at different time (Fig.3A). The comparison of angiogenic capabilities among four groups was calculated at 24 hours and 48 hours, respectively, (Fig.3B-C). Results were expressed as the mean ± S.E. (n=3 per group). \*\**p*<0.01 and \**p*<0.05.

### Lenti-synd4 shRNA inhibited PKCα and ERK1/2 signal pathways in HUVECs cultured with YKL-40

Activation of PKCα and ERK1/2 is regarded as one of the main signaling pathways for migration and proliferation of endothelial cells. Therefore, we investigated the effects of YKL-40 on the expression of PKCα and ratio of p-ERK1/2 to ERK1/2. The Western blot analysis showed that after being rectified by control, the levels of PKCα (1.38±0.02 vs 1.07±0.02, p<0.001) and relative value of p-ERK1/2 to ERK1/2 (1.20±0.01 vs 0.81±0.01, p<0.001) between YKL-40 group and YKL-40+lenti-synd4 shRNA group were significantly different at 24 hours (Fig.4A, 4C and 4D). As expected, the protein levels of PKCα (1.92±0.03 vs 1.45±0.04, p<0.001) and p-ERK1/2 to ERK1/2 (1.36±0.02 vs 0.91±0.01, p<0.001) at 48 hours were also higher in YKL-40 group than those in YKL-40 combined with knockdown of syndecan-4, which were shown in Fig.4B, 4E and 4F.

**Fig. 4.**
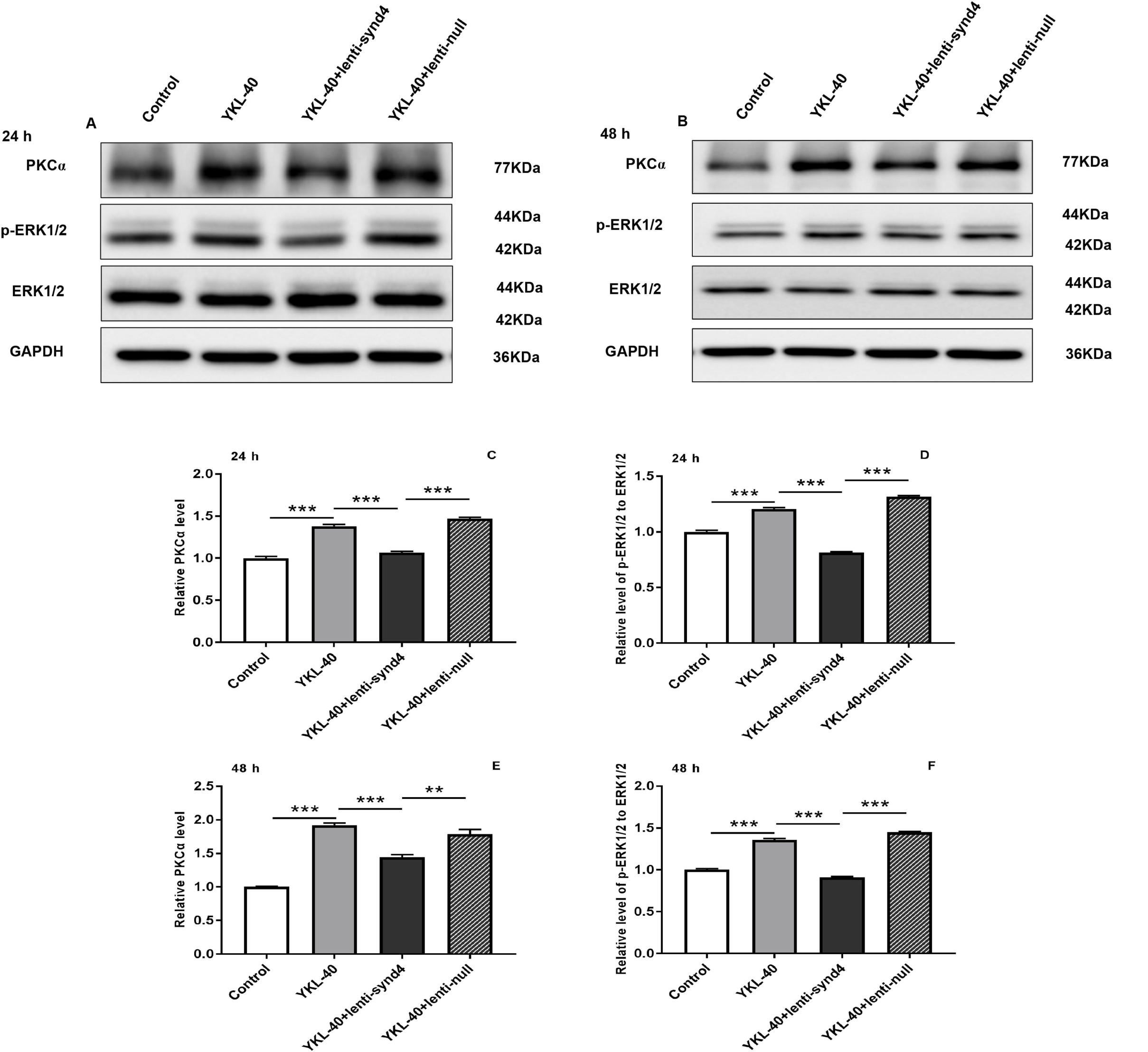
The protein expression level of PKCα and ratio of p-ERK1/2 to ERK1/2 in HUVECs cultured with YKL-40 was inhibited by lenti-synd4 shRNA. The protein expression of PKCα and the ratio of p-ERK1/2 to ERK1/2 in different groups were determined by western blot analysis at 24 hours (Fig. 4A, 4C-D) and 48 hours (Fig.4B, 4E-F). Results were expressed as the mean ± S.E. (n=3 per group). \*\**p*<0.01 and *** *p*<0.001.

## Discussion

Recently, YKL-40 also named chitinase-3-like-1 was considered as a pro-inflammatory and proliferation cytokine in the atherosclerotic lesions.^1,15^ In clinical investigations, Several studies demonstrated that elevated serum YKL-40 levels were independently associated with the presence and extent of coronary artery disease and even higher YKL-40 levels were observed in patients with myocardial infarction.^5,6^ Moreover, increased concentration of serum YKL-40 was closely related to all-cause as well as cardiovascular mortality in an elderly population.^7^ The above studies showed that YKL-40 was possibly associated with unstable plaques, however, the concrete mechanism of YKL-40 on atherosclerosis was still unclear. In our present study, we demonstrated that YKL-40 promoted migration and tube formation of HUVECs, which was partly mediated by syndecan-4.

Increasing evidence suggests that angiogenesis in the atherosclerotic lesions is an important contributors to unstable plaques.^3,16^ The newly formed microvessels are often presented with immaturity and high permeability, permitting inflammatory cytokines to infiltrate into the atherosclerotic plaques; On the other hand, inflammatory state is a strong inducer of angiogenesis as it promotes the release of inflammatory cytokines which facilitate the initiation and development of angiogenesis.^17, 18^ It was reported that YKL-40 modulated vascular endothelial cell morphology by promoting the formation of branching tubules, which showed that YKL-40 have a role in promoting angiogenesis.^19^ Recently, it has been extensively demonstrated that YKL-40 accelerated tumor angiogenesis and was a potential modulator of inflammatory tumor microenvironment.^20,21^ The effects of YKL-40 on angiogenesis involved several signaling pathways, such as focal adhesion kinase (FAK)/protein kinase B (AKT) mentioned in various studies.^22,23^ Another study indicated that signaling pathway of mitogen-activated protein kinase (MAPK) also played a critical role in angiogenesis stimulated by YKL-40, and syndecan-1 may be as one of the membrane receptors for YKL-40.^13^

Syndecans are a small family of four transmembrane proteoglycans, and they have a similar molecular construction including an N-terminal ectodomain, single transmembrane locus and C-terminal cytoplasmic domain.^24^ The importance of syndecans is highlighted by an ability to interact with a series of ligands, including extracellular matrix glycoproteins, growth factors, morphogens, and cytokines which are important regulators of regeneration.^10^ Syndecan-4 is one of the members of syndecans, which are endowed with multiple role including migration, proliferation and homeostasis.^25^ Unlike other syndecans, syndecan-4 has a widespread tissue distribution, which is abundantly expressed in endothelial cells, and it also plays an important role in the development of angiogenesis.^26^ Delivery exogenous syndecan-4 could improve angiogenic therapy for ischemia in diabetes; on the contrary, decreasing the expression of syndecan-4 on the cell surface remarkably inhibited the adhesion and migration of endothelial progenitor cells.^27,28^ In the present study, we found that YKL-40 prompted the migration and tube formation of HUVECs, and knockdown of the syndecan-4 inhibited angiogenic potential of HUVECs stimulated by YKL-40. Hence, our study showed that syndecan-4 was possibly a mediator for YKL-40-induced angiogenesis.

Protein kinase C (PKC) and extracellular regulated protein kinases 1/2 (ERK1/2) are main signal pathways involved in the regulation of proliferation, differentiation, cell migration, adhesion and apoptosis.^29-31^ It has been shown that the syndecan-4 KKXXXKK motif of V region combines with phosphotidylinositol (4,5)-bisphosphate and promotes the stabilization of syndecan-4 dimers.^32,33^ Moreover, this motif is an important bonding location for protein kinase C-alpha (PKCα). Syndecan-4 interacts with PKCα and this interaction regulates localization of PKCα into the cytoskeleton, resulting in a sustained PKC activity.^34^ As one of mitogen-activated protein kinase (MAPK) family members, the activation of ERK1/2 has also been reported in the angiogenesis induced by syndecan-4.^35^ In the current study, we further demonstrated that YKL-40 promoted the angiogenesis associated with the activation of PKCα and ERK1/2 signal pathways, and silencing the gene production of syndecan-4 in HUVECs with lenti-synd4 shRNA significantly down-regulated the expression of PKCα and ratio of p-ERK1/2 to ERK.

In conclusion, our study indicated that the effects of YKL-40 on migration and tube formation of HUVECs were partly regulated by syndecan-4 through PKCα and ERK1/2 signal pathways. It is also clear that YKL-40 is a potential cytokine in promoting angiogenesis and is regarded as a prognostic factor for cardiovascular diseases in previous studies. However, we could not make a definite conclusion that YKL-40 was associated to vulnerable atherosclerotic plaque formation and plaque rupture according to current study. In order to verify the hypothesis of YKL-40 promoting unstable atherosclerotic plaque, constructing suitable animal experiments to clarify this mechanism is indispensable in the following studies.

## Author’s Contributions

WeiJun Sun and Jianlei Zheng designed the study. Jianei Zheng and Qi Xue performed the experiments, and the data were also analyzed and interpreted by Jianei Zheng and Qi Xue. Yan Zhao wrote the manuscript. All authors read and approved the final manuscript.

